# Immunomodulatory effects of extract of *Ganoderma lucidum* basidiocarps cultivated on alternative substrate

**DOI:** 10.1101/603498

**Authors:** Biljana Božić Nedeljković, Jasmina Ćilerdžić, Dragana Zmijanjac, Milan Marković, Tanja Džopalić, Saša Vasilijić, Mirjana Stajić, Dragana Vučević

**Affiliations:** Institute for Medical Research, Military Medical Academy, Belgrade, Serbia; University of Belgrade, Faculty of Biology, Serbia; University of Nis, Faculty of Medicine, Serbia; Faculty of Medicine of the Military Medical Academy, University of Defence in Belgrade, Serbia

**Keywords:** Antigen presenting cell, *Ganoderma lucidum*, Wheat straw, Immunomodulation

## Abstract

The aim of the study was to investigate if there are any differences in effects of extracts of commercially (GC) and alternatively (wheat straw) (GA) cultivated *Ganoderma lucidum* basidiocarps on properties of peritoneal macrophages (PM) and monocyte-derived dendritic cells (MoDCs). GA extract stronger stimulated the metabolic and phagocytic activity of PMs, their adhesion capability and ability to produce ROS and NO compared to GC. Both tested extracts significantly increased allostimulatory and Th1 polarization capacity of simultaneous TLR3 and TLR7-activated MoDCs, but GA extract was more effective. The GA extract increased the production of ROS and NO by TLR4 stimulated PMs and up-regulated the production of certain cytokines as well as allostimulatory and Th1 polarization capacity of MoDCs. The GA extract could be a potent immunostimulatory agent for activation of MoDCs with the simultaneous engagement of TLRs that seems to be a promising strategy for preparation of DC-based anti-tumor vaccines.

## Introduction

*Ganoderma lucidum* (Curtis) P. Karst., known as Reishi, is a popular medicinal mushroom used in traditional medicine for the prevention and treatment of various pathological conditions. Today, there is evidence that bioactive constituents of this species are responsible for numerous health benefits due to strong immunomodulatory, antitumor, antioxidative, anti-inflammatory, antimicrobial, and many other activities (Lin, Kao et al. 2006, Yuen and Gohel 2008, Joseph, Sabulal et al. 2011, Wasser 2014). Therefore, commercial production of this mushroom is continuously growing and nowadays the emphasis is put on the creation of a system for cheaper, easier, faster and environmentally friendly cultivation of biologically more active fruiting bodies. Thus, fruiting bodies with higher immunostimulatory potential could be applied as natural pharmaceutical agents in immunotherapy of patients suffering from various tumor types (Park, Kuen et al. 2018).

The antigen-presenting cells (APCs) are a common checkpoint for stimulation of the immune system and induction of potent antitumor response in cancer treatment (Martin, Schreiner et al. 2015). They include dendritic cells (DCs), macrophages, and B lymphocytes, which participate in capturing, processing and presenting antigens to T lymphocytes (Constantino, Gomes et al. 2017). Macrophages and DCs are the powerful phagocytic cells, key players in the innate immune system and link between innate and adaptive immunity, which are derived from peripheral blood monocytes and exist in almost all tissues (Clark, Angel et al. 2000, Hirayama and Iida 2017). Activated macrophages produce numerous bioactive compounds, including reactive oxygen species (ROS), nitric oxide (NO), an important mediator of innate immune response on various pathological stages, as well as cytokines, primarily interleukins (IL-1, IL-6), tumor necrosis factor α (TNF-α), and interferon-γ (IFN-γ), which are crucial in recruitment and activation of other immune cells and stimulation of adaptive immunity (Hirayama and Iida 2017). In the presence of microbes or inflammatory stimuli, DCs undergo a complex process of maturation that includes up-regulation of co-stimulatory molecules, migration to lymph nodes, T lymphocytes priming and cytokine production (Clark, Angel et al. 2000). These potent APCs express various pattern recognition receptors (PRRs) and in such a way trigger signalling pathways resulting in their phenotypic changes and functional maturation (Martin, Schreiner et al. 2015). Toll-like receptors (TLRs) present an important group of PRRs on the macrophages and DCs surface and crucial factors for recognition of viruses, bacteria, fungi, and parasites, i.e. they play a key role in innate immunity (Kawai and Akira 2009). These authors emphasized that ligation of different TLRs by specific TLR agonists presents a powerful tool for induction of DCs maturation. TLR agonists are used as adjuvants or immune modifiers in DC-based trials of tumor immunotherapy) (Gnjatic, Sawhney et al. 2010). However, since single TLR agonist has relatively limited adjuvant effects on DC phenotype and function, current studies are focused on research of synergy between paired TLR agonists (Zheng, Cohen et al. 2008).

Starting from the fact emphasized by Pi and colleagues (Pi, Chu et al. 2014) that polysaccharides of *Ganoderma* spp. possess strong immunostimulatory activity based on their recognition as foreign molecules by various PRRs on DCs, and consequently, on stimulation of APCs maturation, the aim of this study was defined. In our study, we investigated the potential synergism of extracts of commercially (GC) and alternatively (wheat straw) (GA) cultivated *G. lucidum* basidiocarps with different TLRs on different APCs. Namely, we investigated the immunomodulatory effects of GC and GA extracts on functional properties of peritoneal macrophages stimulated by TLR4 and on functional characteristics of human monocyte-derived dendritic cells stimulated by simultaneous engagement of TLR3 and TLR7.

## Material and methods

### Organism and growth conditions

The culture of *Ganoderma lucidum* BEOFB 431, isolated from fruiting body collected in Bojčin forest (Belgrade, Serbia), is maintained on Malt agar medium in the culture collection of the Institute of Botany, Faculty of Biology University of Belgrade. The fruiting bodies were cultivated on alternative substrate consisted of wheat straw under laboratory conditions. Basidiocarps of a commercial Chinese strain, cultivated on oak sawdust, were purchased at a health food store.

### Preparation of the basidiocarp extracts

The dried and pulverized commercially and alternatively produced *G. lucidum* basidiocarps (2.0 g) were extracted with 60.0 ml of 96% ethanol by stirring on a magnetic stirrer (150 rpm) for 72 h. The resultant extracts were centrifuged (20 °C, 3000 rpm, 10 min) and supernatants were filtered through Whatman No.4 filter paper, concentrated under reduced pressure in a rotary evaporator (Büchi, Rotavapor R-114, Germany) at 40 °C to dryness, and dissolved in 5% dimethyl sulphoxide (DMSO) to an initial concentration of 10.0 mg/ml.

### Experimental animals

All animal experiments were approved by the Ethics Review Committee for Animal Experimentation of Military Medical Academy and Ministry of Agriculture and Environmental Protection of Republic of Serbia (Veterinary Directorate No. 323-07-7363/2014-05/5). Inbred male Albino Oxford rats (AO; *Vivarium* for Small Experimental Animals, Military Medical Academy, Belgrade) weight about 200-220 g were housed in an air-conditioned room at 25 °C on a 12h-light/dark cycle. Animals were provided pelleted food (Veterinary Institute, Subotica) and tap water *ad libitum.* Sacrifice was done with intravenous injection of Ketamin/Xilazyn in a lethal dose. All procedures were done in accordance with the Guide for the care and use of laboratory animals.

### Peritoneal macrophages isolation and experimental design

The medium used for the cell isolation and incubation was HEPES-buffered Roswell Park Memorial Institute medium (RPMI-1640) supplemented with fetal calf serum (FCS) (Flow, Irvine, Ca, USA), glutamine (ICN Flow, SAD), penicillin, and gentamicin (Galenika a.d.d., Serbia).

Peritoneal cells were obtained by sterile lavage with RPMI medium supplemented with 2% FCS and heparin (Galenika a.d.d., Serbia). Enrichment of peritoneal cell exudates with PMs was enabled using density gradient OPTIPREP (Nycomed Pharmas, Norway) with 0.8% NaCl. After centrifugation on the gradient, mononuclear cells (highly enriched with PM, >90%) were washed and resuspended in RPMI-1640 supplemented with 10% FCS and cell number was adjusted to 10^6^ cells/ml. Afterwards, the cells were seeded in 96-well plate in two ways: *i*) 1×10^5^ cells per well for testing the viability and production of phagocytic activity, ROS, NO, and cytokine and *ii*) 5×10^5^ cells per well for assessment of adhesion capacity.

Peritoneal macrophages (PMs) isolated in this way were cultivated under standard conditions (37 °C, 5% CO_2_) for 24 h, and treated with GC and GA extracts in final concentration of 100.0, 10.0, and 0.1 μg/ml per well in presence or absence of adequate stimulator (depending of evaluated function). Lipopolysaccharide (LPS, Sigma, USA), TLR4 agonist, at a final concentration of 100.0 ng/ml per well, was used as a stimulator for assessment of metabolic viability, phagocytic activity, and NO production. Adhesion capacity and ROS production were assessed by phorbol-myristate-acetate (PMA, Sigma, USA) at the final concentration of 250.0 ng/ml per well. Control cells were cultivated under standard conditions, with or without TLR4 agonist and were not treated with GC and GA extracts. All studied functions of PMs were observed after 24 h cultivation *in vitro* and were done in quadruplicate.

### Cell viability assay

Cell viability was estimated by a quantitative colorimetric assay described for human granulocytes which were based on metabolic reduction of 3-(4,5-dimethylthiazol-2-yl)-2,5-diphenyltetrazolium bromide (MTT, Invitrogen) into coloured product formazan (Oez, Platzer et al. 1990). MTT assay was conducted with 24 h cultivated PMs and MTT which was added in the concentration of 5.0 mg/ml (10.0 μl per well), which were incubated at 37 °C in an atmosphere of 5% CO_2_ and 95% humidity for 3 h. The absorbance of produced formazan after overnight incubation in the solution composed of sodium dodecyl sulphate (SDS) and HCl (10% SDS with 0.01 N HCl) was measured at dual wavelengths, 570/650 nm by an ELISA 96-well plate reader (Behringer, Germany). Cells viability was expressed as absorbance of solubilized formazan at the end of the incubation period. The presented data are expressed as the mean ± standard error (SE) from three independent experiments performed in triplicate.

### Phagocytosis assay

The phagocytic capacity of PMs was determined according to the technique described by Chen and colleagues (Chen, Weng et al. 2015). After 24 h cultivation PMs without/with stimulators and GC/GA extract, the supernatants were collected and 50 μl/well of neutral red (1:300) was added and incubated for 4 h. After incubation, supernatants were discarded, cells were washed with phosphate-buffered saline (PBS) three times and lysed by adding 100.0 μl/well of cell lysing solution (ethanol and 1% acetic acid at the ratio of 1:1), and absorbance of the solution was measured at 540/650 nm using Microplate Reader (Behringer, Germany). The presented results are expressed as the mean ± SE from three independent experiments performed in triplicate.

### Adhesion capacity assay

Adhesion capacity of PMs was assessed by a method of Oez and colleagues (Oez, Welte et al. 1990) based on the cell ability to adhere to the plastic matrix. After the 24 h cultivation without/with stimulator (PMA) and GC/GA extract, supernatants were removed and cells were washed three times with warm PBS in order to remove non-adhered cells. Then in each well added methanol (100.0 μl/well) and it was incubated for seven minutes. Attached cells were dyed with 0.1% solution of methyl blue (100.0 μl/well) for 15 minutes and washed three times with tap water. Plates were left to dry on air overnight and colour was dissolved by adding 0.1 N HCl (200.0 μl/well). The absorbance of the solution in each well was measured at 650/570 nm using Microplate Reader (Behringer, Germany). The presented results are expressed as the mean ± SE from three independent experiments performed in triplicate.

### NBT reduction assay

NBT assay was used to evaluate the generation of superoxide anion (O_2_^-^) produced by PMs (Pick, Charon et al. 1981). Briefly, nitroblue tetrazolium (NBT, Invitrogen), in final concentration of 0.5 mg/ml per well, was added to PMs suspension after 24 h treatment of PMs without/with stimulators and GC/GA extracts and the mixture was incubated at 37 °C in an atmosphere of 5% CO_2_ and 95% humidity for one hour. Formed diformazan crystals were dissolved by adding SDS-HCl mixture (100.0 μl/well) and optical density was measured at 570/650 nm by a Microplate reader (Behringer, Germany). The presented results are expressed as the mean ± SE from three independent experiments performed in triplicate.

### Determination of NO production

Production of NO was quantified by the accumulation of nitrite as a stable end-product and determined by a Greiss reaction assay (Green, Wagner et al. 1982). Equal volumes of the supernatants and Griess reagent [0.35% 4-aminophenyl sulfone (Sigma-Aldrich, Germany), 0.1% N-(1-naphthyl)ethylenediamine dihydrochloride in 1M HCl (POCh, Poland)] were incubated at room temperature (22 ± 2 °C) for 10 minutes. The optical density of the solution was measured at 540/650 nm using Microplate Reader (Behringer, Germany). The nitrite concentration (μM) was calculated from the prepared standard curve for the known NaNO2 concentrations. The presented results are expressed as the mean ± SE from three independent experiments performed in triplicate.

### Preparation and treatment of human monocyte-derived dendritic cells

Immature monocyte-derived dendritic cells (MoDCs) were generated from the adherent fraction of human peripheral blood mononuclear cells (PBMCs). Namely, PBMCs from buffy coats of healthy volunteers (upon written informed consent) were isolated by density centrifugation in Lymphoprep (Nycomed, Oslo, Norway), re-suspended in 5.0 ml of 10% FCS with 2-Mercaptoethanol (2-ME) in RPMI medium and allowed to adhere to plastic flasks. After incubation at 37 °C for 90 minutes, non-adhered cells were removed and adhered cells were cultured in 5.0 ml of RPMI medium containing granulocyte-macrophage colony-stimulating factor (GM-CSF; 100.0 ng/ml) and IL-4 (20.0 ng/ml). On day three, half of medium volume was removed and replaced with the same volume of fresh medium containing GM-CSF and IL-4 and it was incubated for next two days. At the end of the incubation period (on day five), MoDCs (5×10^5^ cells/well) were moved in 24-well plate, in RPMI medium containing GM-CSF/IL-4. After six days, immature MoDCs were replated (5 x 10^5^ cells/ml) in medium with different combination of TLR3 agonist (Poly (I:C), Sigma-Aldrich, Germany, 10.0 μg/ml) and TLR7 agonist (Loxoribine Sigma-Aldrich, Germany, 34.0 μg/ml) with GC and GA extracts (100.0 μg/ml), and incubated for 24 h. Afterwards, cell-free supernatants were collected for cytokine analysis, while cells were detached and their phenotype was observed. Cell-free supernatants were collected and stored at −20 °C for the subsequent determination of cytokine levels. The cells were used for further studies.

### Flow cytometry analysis of MoDCs for immunophenotyping

MoDC were obtained by cultivation of human monocytes for six days with GM-CSF (100.0 ng/ml) and IL-4 (20.0 ng/ml). After this period MoDC additionally cultivated in presence of Poly (I:C) (TLR3 agonist) (10.0 μg/ml) and Loxoribine (TLR7 agonist) (34.0 μg/ml) and GC or GA extracts (100.0 μg/ml). Control (non-treated) and those treated MoDCs (2×10^5^ cells/tube) were washed in PBS supplemented with 2% FCS and 0.1% NaN3, and incubated at 4 °C for 45 min with one of the following monoclonal antibodies (mAbs): HLA-DR coupled with phycoerythrin (PE), CD83 conjugated with fluorescein isothyocianate (FITC), CD86-PE, CD40-FITC, CD54-PE (Serotec, Oxford, UK), and CCR7-FITC (R&D Systems, Minneapolis, MN, USA). The control MoDCs incubated with irrelevant mouse Mab reactive with rat antigens were used as the irrelevant control. The cells were incubated at 4 °C for 45 min, washed and fixed with 1% paraformaldehyde. For cell fluorescence analysis, at least 5×10^3^ cells per sample were analysed using EPICS XL-MCL flow cytometer (Coulter, Krefeld, Germany). The cell-surface expression on MoDCs was determined by means of a forward versus side scatter gate.

### Mixed leukocyte reaction with allogeneic CD4^+^ T lymphocytes

Conventional CD4^+^ T lymphocytes were isolated from PBMCs using negative immunomagnetic sorting with CD4^+^ T lymphocytes isolation MACS kit (MyltenyiBiotec, Germany). According to flow cytometry analysis, the purity of isolated CD4^+^ T lymphocytes was higher than 95%. Purified allogeneic CD4^+^ T lymphocytes (10^5^ cells/well) were placed in 96-well plates and cultivated with MoDCs pre-treated with TLR agonists and *G. lucidum* extracts in RPMI medium with 10% FCS. Cell proliferation was detected after five days of cultivation. Cells were pulsed with [3^H^] thymidine (1.0 μCi/well; Amersham, UK) for the last 18 h of cultivation, then harvested onto glass fibre filters, and [3^H^] thymidine incorporation into DNA was measured by β-scintillation counting (LKB-1219; Rackbeta, Finland). Results were expressed as counts per minute (c.p.m.). Values are given as mean ± standard deviation (SD). The presented results are representative of three independent experiments (n = 3 different donors).

### Evaluation of cytokine production

The levels of TNF-α, IL-12, IL-6, and IL-10 were measured in the cell-free supernatants of control and treated MoDCs cultures (4×10^5^ cells/ml) by ELISA assays (R&D Systems, Minneapolis, USA). The levels of cytokines produced by CD4^+^ T lymphocytes in co-cultures with MoDCs were evaluated in the cell-free supernatants of co-cultures. Briefly, purified allogeneic CD4^+^ T cells (1×10^5^ cells/well) were cultivated with MoDCs (1×10^4^ cells/well) pre-treated with TLR agonists and *G.lucidum* extracts in RPMI medium with 10% FCS in 96-well plates. Phorbol-myristate-acetate (20.0 ng/ml) and ionomycin (500.0 ng/ml) (Merck, Austria) were added to the wells after five days. Cells were incubated for an additional 8 h and then harvested, and cell-free supernatants were collected after centrifugation and stored at −20 °C for the subsequent determination of the studied cytokine levels. Values are given as mean ± SD. The presented results are representative of three independent experiments (n = 3 different donors).

### Statistical analysis

Data were analysed for significant differences using Student’s paired *t*-test. A *p* value less than 0.05 was considered to be statistically significant.

## Results

### Effect of Ganoderma lucidum extracts on metabolic activity/viability of PMs

The PMs cultivated with three different concentrations of commercially or alternatively cultivated *G. lucidum* basidiocarps (GC and GA, respectively) (0.1 μg/ml, 10 μg/ml, and 100 μg/ml) without (non-stimulated) (Fig. 1A) or with LPS (1 μg/ml) (LPS-stimulated) (Fig. 1B) during 24 h.

**Figure 1.**
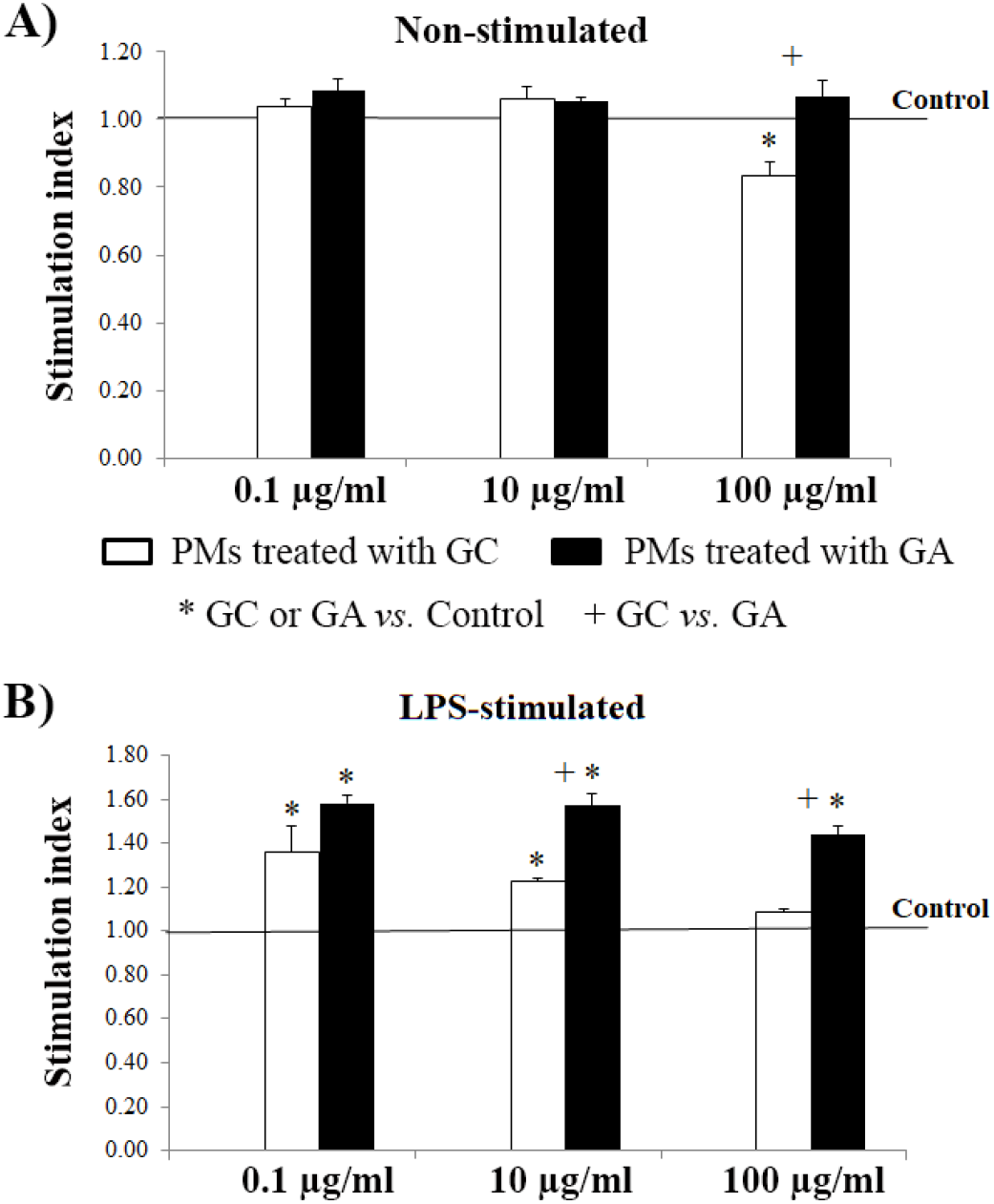
Effect of *Ganoderma lucidum* extracts on metabolic activity/viability of PMs.

The GC and GA extracts have no effect on metabolic activity/viability of non-stimulated PMs, except GC extract in the concentration of 100.0 μg/ml which affected inhibitory (Fig. 1A). However, all tested extracts (except GC in a concentration of 100.0 μg/ml) showed a stimulatory effect on metabolic activity/viability of LPS-stimulated PMs. Furthermore, in comparison with GC, GA has a stronger effect in concentration 10 μg/mL and 100 μg/ml on metabolic activity/viability of LPS-stimulated PMs (Fig. 1B).

### Effect of Ganoderma lucidum extracts on adhesive capability and phagocytic activity of PMs

Effect of *G. lucidum* extracts cultivated on different substrates (commercially and alternatively) on adhesion capacity of non-stimulated and PMA-stimulated PMs are shown in Fig. 2. The PMs cultivated with three different concentrations of GC or GA extracts (0.1 μg/ml, 10 μg/ml, and 100 μg/ml) without (non-stimulated) (Fig. 2A) or with PMA (250 μg/ml) (PMA-stimulated) (Fig. 2B) during 24 h. Adhesive capability of attached PMs was assessed by dyed with methyl blue as described in the Materials and Methods. Treatment of non-stimulated PMs with GC extract did not modulate their adhesive capability, while treatment with GA extract, in all tested concentrations, enhanced their adhesive function compared to the ability of non-treated and GC treated PMs (Fig. 2A). Comparing with a control, GC extract had no effect on the adhesive capacity of PMA-stimulated PMs, while GA extract (in all tested concentrations) enhanced this ability (Fig. 2B).

**Figure 2.**
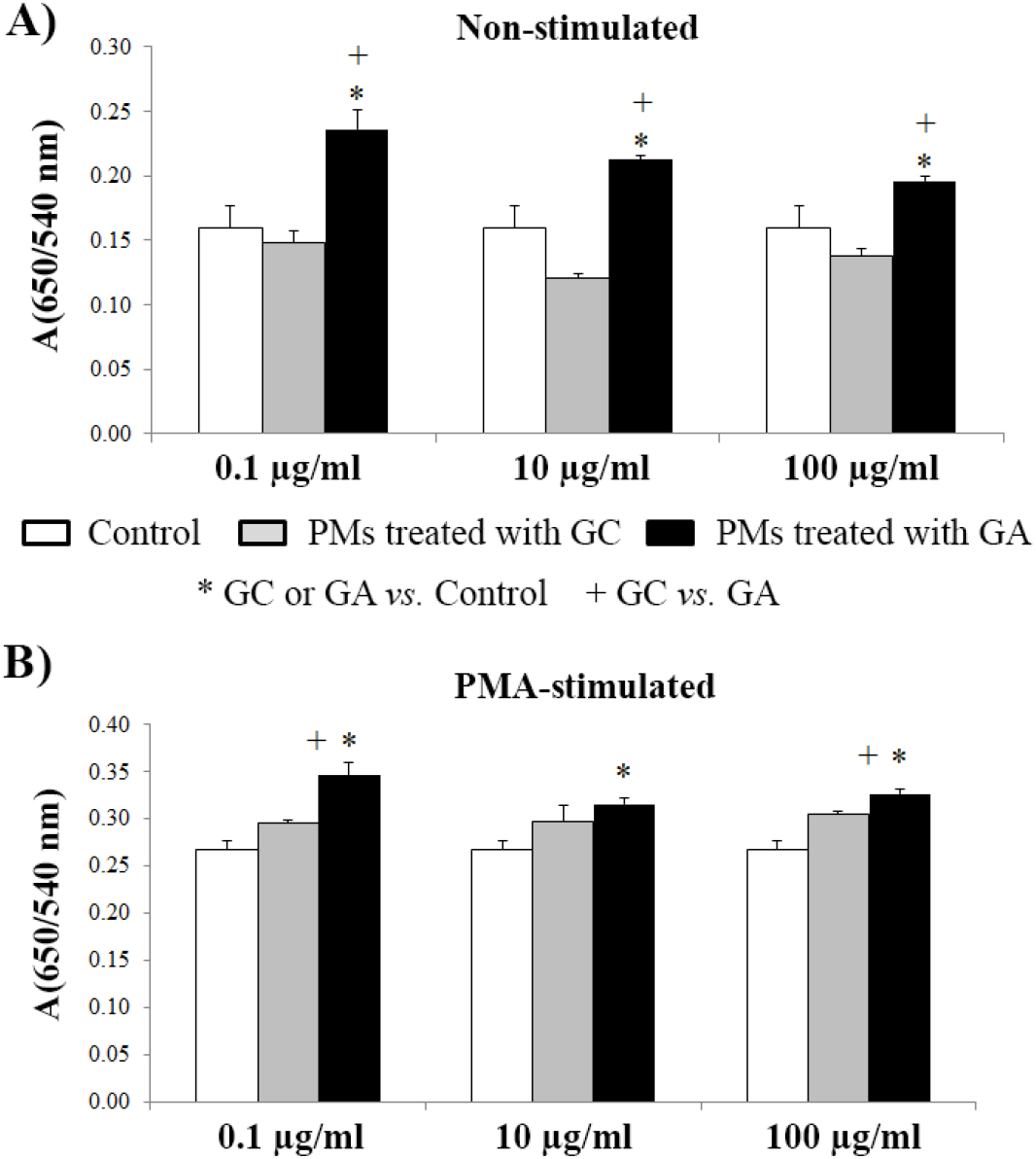
Effect of *Ganoderma lucidum* extracts on adhesive capability of PMs.

The PMs cultivated with three different concentrations of GC or GA extracts (0.1 μg/ml, 10 μg/ml, and 100 μg/ml) with LPS (1 μg/ml) (LPS-stimulated) during 24 h (Fig. 3). Phagocytotic/pinocityc activity was assessed by Neutral red uptake as described in the Materials and Methods section.

**Figure 3.**
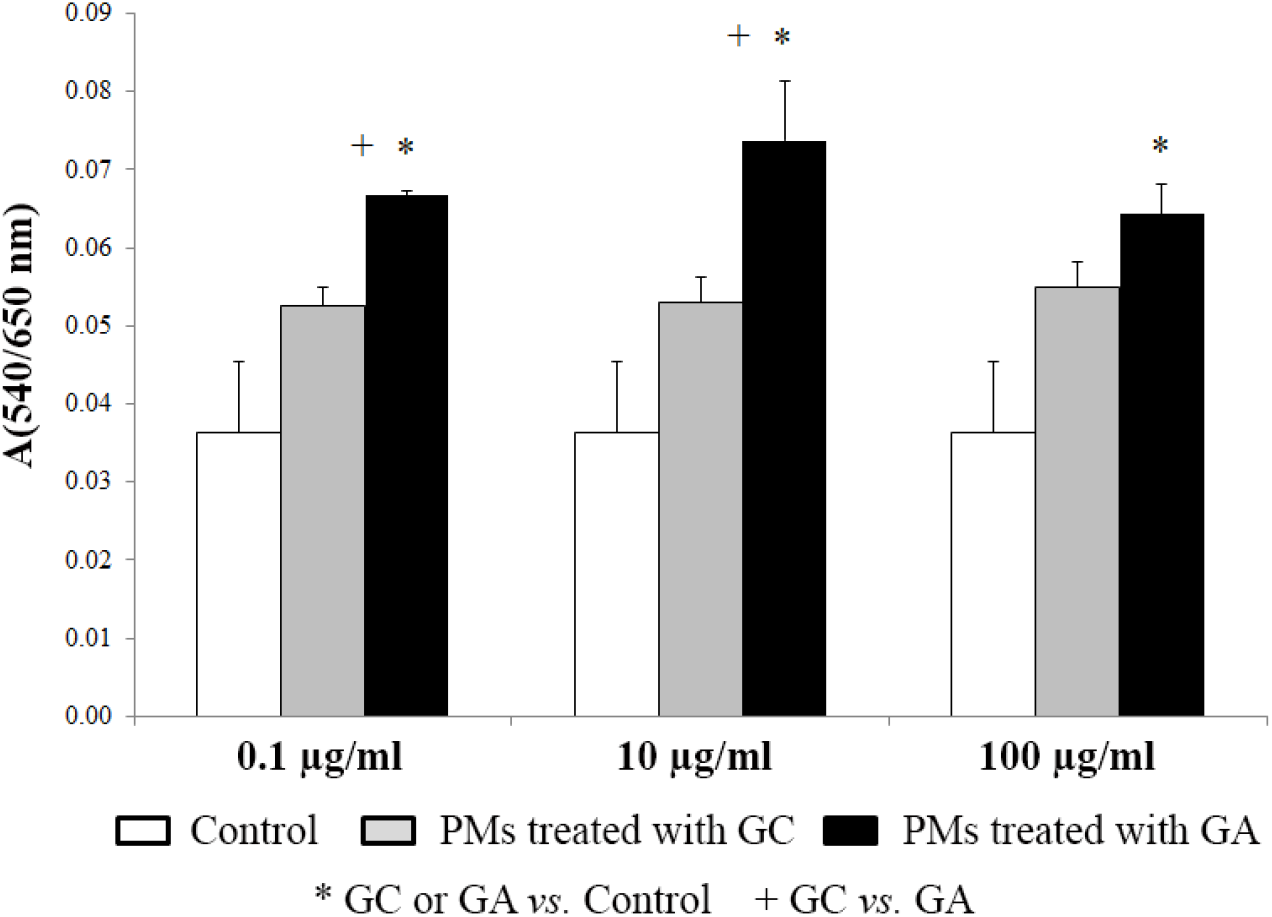
Effect of *Ganoderma lucidum* extracts on phagocytic activity of LPS stimulated PMs.

GC and GA extracts showed no significant effect on the phagocytic function of nonstimulated PMs (data not shown). However, in the case of LPS-stimulated PMs, GA extracts (in all tested concentrations) statistically considerable up-regulated this activity in comparison with nontreated PMs and PMs treated with GC extract (Fig. 3).

### Effect of Ganoderma lucidum extracts on ROS and NO production by PMs

Effect of investigated extracts on the ROS production by PMs treated with LPS after 24 h treatment compared to the control cell (LPS-stimulated) measured by NBT test as described in the Materials and Methods (Fig. 4). The treatment of non-stimulated PMs with GC and GA extracts did not influence their potential of ROS production (data not shown). Comparing with adequate control cells, GC extract had no effect on ROS production by LPS-stimulated PMs, contrary to GA extract which induced a statistically significant increasing of this ability (Fig. 4).

**Figure 4.**
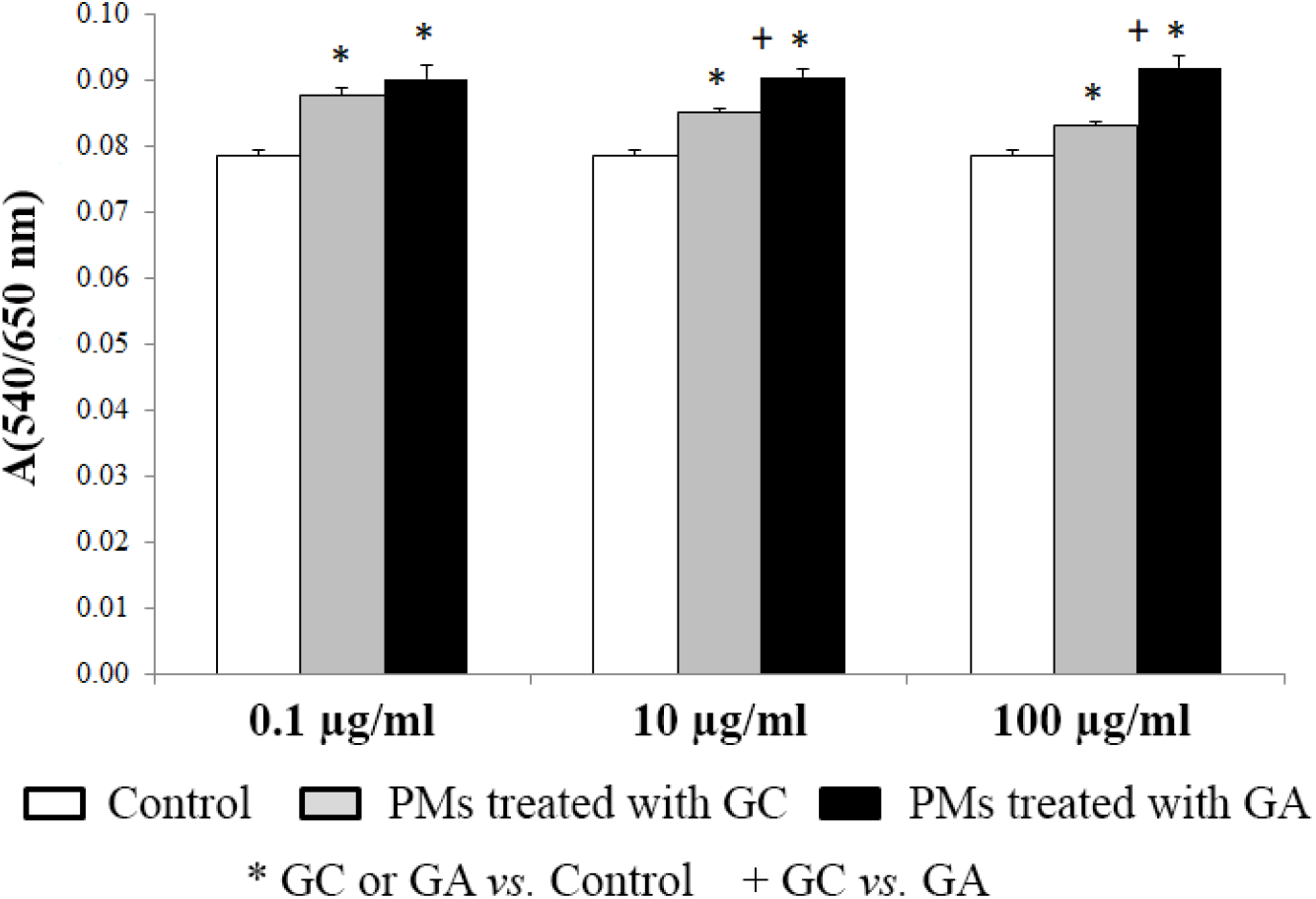
Effect of *Ganoderma lucidum* extracts on ROS production by LPS stimulated PMs.

Effect of investigated extracts on the nitrite level in supernatants of PMs treated with/without LPS after 24 h treatment compared to control cell (non-stimulated or LPS-stimulated) measured by Griess test as described in the Materials and Methods (Fig. 5). The treatment of nonstimulated and LPS-stimulated PMs with GC and GA extracts, in all tested concentrations (except treatment of non-stimulated cells with 100.0 μg/ml of GC extract) caused an increase of NO production. Furthermore, GA extract was a statistically stronger inducer of NO production by tested PMs than GC extract (Fig. 5A and 5B).

**Figure 5.**
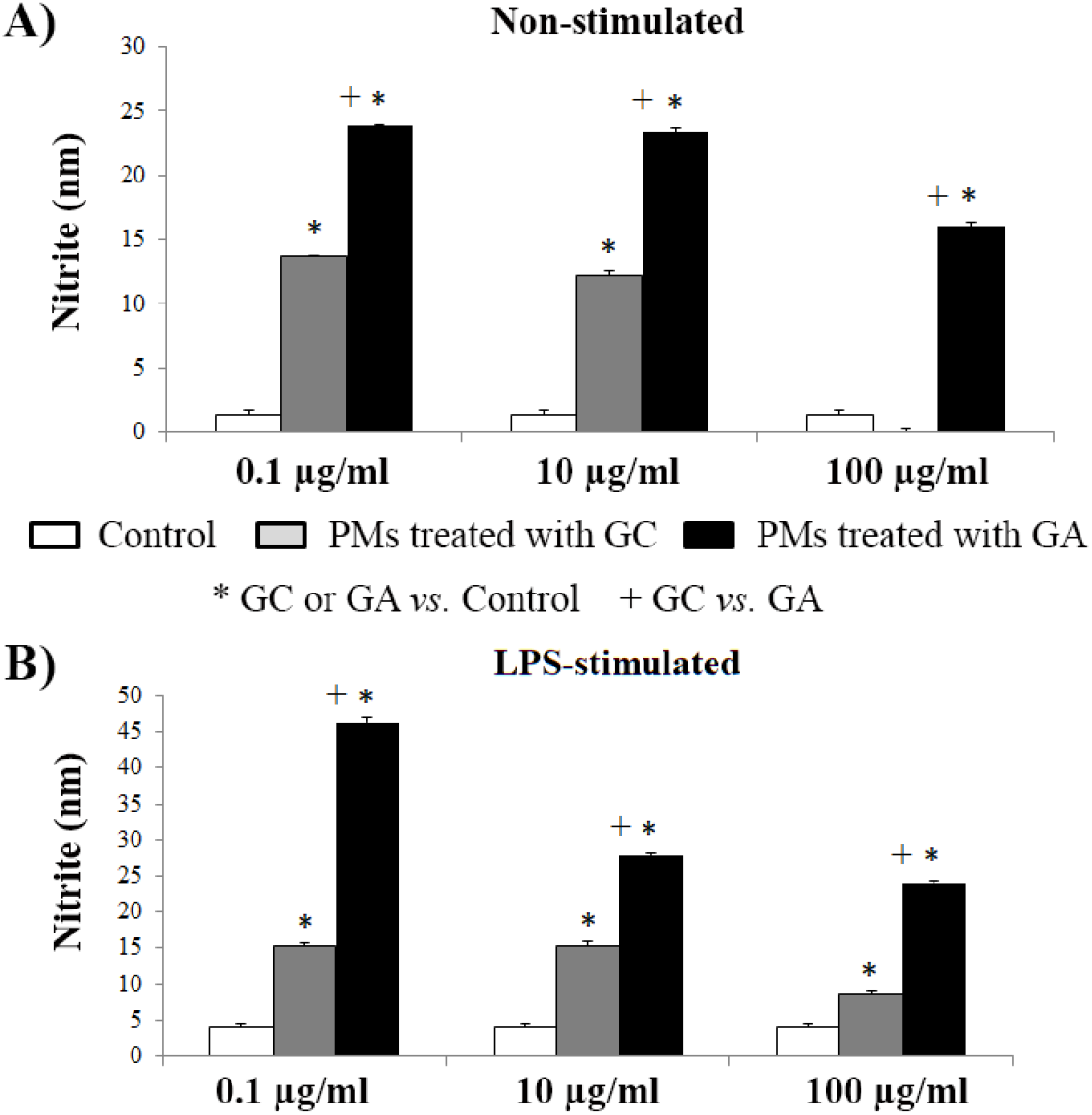
Effect of *Ganoderma lucidum* extracts on NO production by PMs.

### Effect of Ganoderma lucidum extracts on phenotype and cytokine production by MoDCs

Phenotype analysis showed that GC and GA extracts did not affect MoDCs phenotype measured by percentage of HLA-DR, CD83, CD86, CD40, and CCR7 positive cells (data not shown). Simultaneous treatment of MoDCs with treatment with Poly (I:C), Loxoribine and GC or GA extract up-regulated mean of fluorescence (MnI) of CD83 and HLA-DR. Additionally, simultaneous treatment of TLR3 and TLR7-stimulated MoDCs with GA extract also affected the expression of co-stimulatory molecules, CD86 and CD40, as well as a chemokine receptor, CCR7 (Fig. 6A).

**Figure 6.**
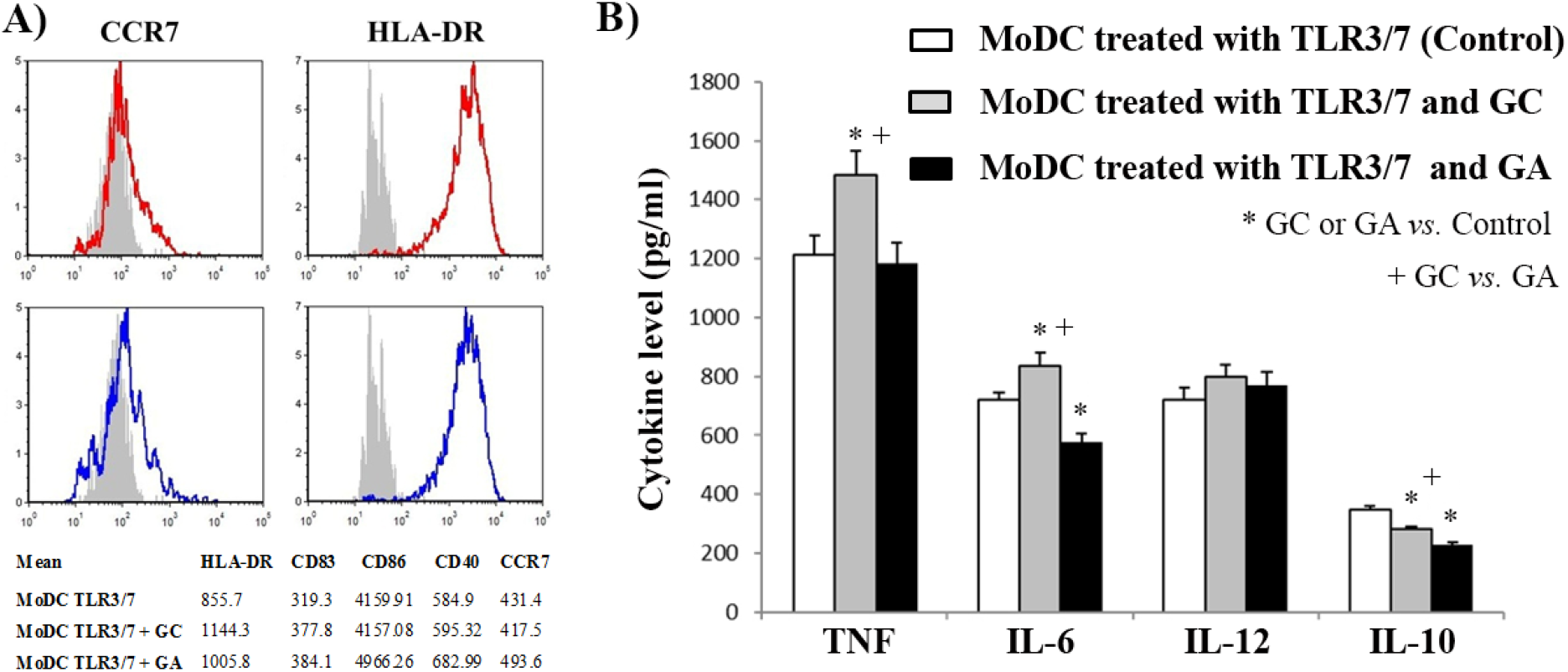
Effect of *Ganoderma lucidum* extracts on phenotype (The red line represents expression for MoDC TLR3/7 + GC, the blue line represents expression for MoDC TLR3/7 + GA, the full gray histogram represents expression for MoDC TLR3/7) (A) and cytokine production (B) by MoDCs.

The additions of GC extract stimulated the production of IL-6 and TNF-α, had no effect on the production of IL-12, and inhibited the production of IL-10 by TLR3 and −7 stimulated MoDCs. On the other hand, GA extract inhibited the production of IL-6 and IL-10 and had no effect on TNF-α and IL-12 production by studied MoDCs compared to MoDC treated only with TLR3 and TLR7 antagonist (Fig. 6B).

### Effect of Ganoderma lucidum extracts on alostimulatory and Th cells polarization capacity of MoDCs

The ability of TLR3 and TLR7-stimulated MoDCs to induce the proliferation of allogeneic CD4^+^ T lymphocytes was not changed by the addition of GC extract, while GA extract significantly enhanced this ability (Fig. 7A). MoDCs treated with TLR3 and TLR7 agonists and with GC extract down-regulated production of IFN-γ and had no effect on IL-17 production by allogeneic CD4^+^ T lymphocytes compared to the effect of MoDCs treated only with TLR3 and TLR7 agonists. On the other side, GA extract in combination with TLR3 and TLR7 changed the properties of MoDCs and thus stimulated MoDCs up-regulated IFN-γ synthesis and had no effect on IL-17 generation by allogeneic CD4^+^ T lymphocytes in comparison with TLR3 and TLR7-stimulated MoDCs (Fig. 7B). Additionally, GA extract in combination with TLR3 and TLR7 up-regulated IL-17 synthesis by allogeneic CD4^+^ T lymphocytes in comparison with GC, TLR3 and TLR7-stimulated MoDCs (Fig. 7B).

**Figure 7.**
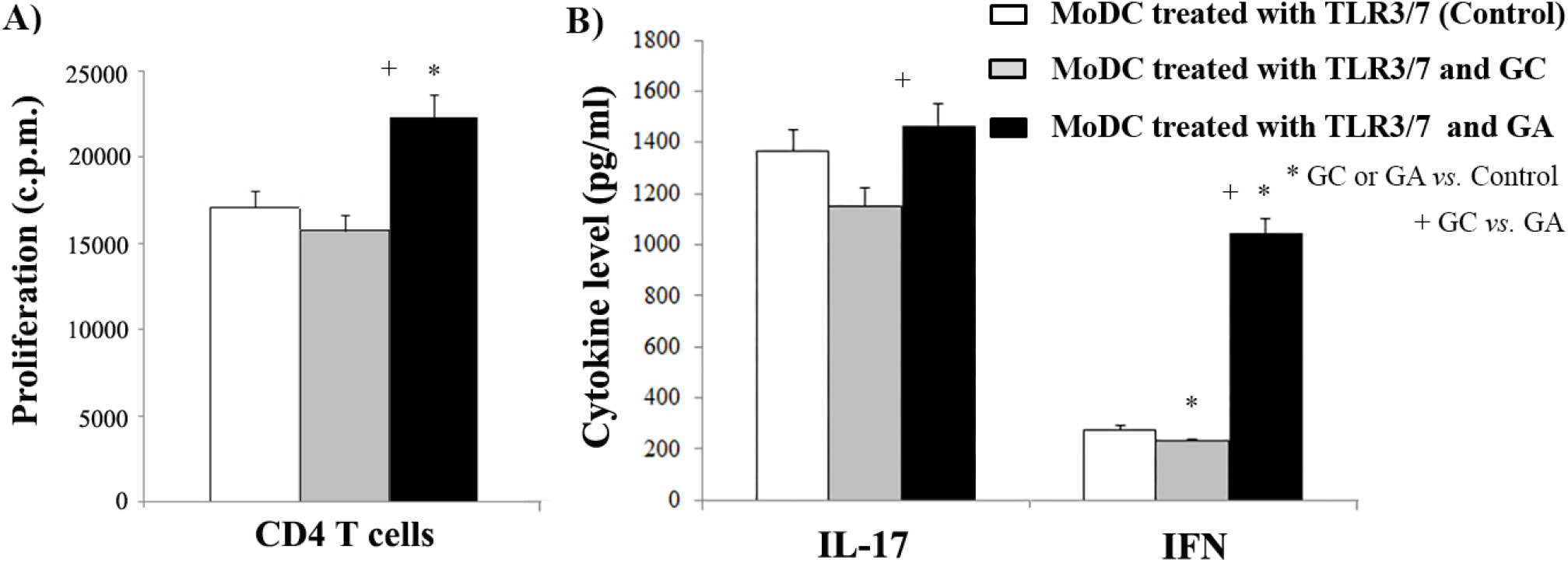
Effect of *Ganoderma lucidum* extracts on alostimulatory (A) and Th polarization capacity (B) of MoDCs

## Discussion

Cooperation of different PRR signals in APC during the induction of immune responses is an emerging field in innate immune research. Activation of two or more TLRs, or other PRRs at the same time, which mimic the actual situation during host cell–microbe interaction, may lead to synergistic, antagonistic, or additive effects(Makela, Strengell et al. 2009). On the other side, *Ganoderma* spp. extracts and metabolites (derived from wild and/or traditionally cultivated basidiocarps) express strong immunomodulatory characteristics and present effective modifiers of different biological processes (Lin, Chen et al. 2006, Boh 2013, Shi, Zhang et al. 2013, Liu, Xing et al. 2016, Liu, Xing et al. 2016). According to available literature, there is no knowledge about the immunomodulatory potential of crude extracts of *G. lucidum* basidiocarps cultivated on any alternative and environmentally-friendly substrate. This is the first report on it and our findings showed that crude extract of *G. lucidum* basidiocarps cultivated on nutritionally poor wheat straw contained moleculeswith higher immunomodulatory potentials that in cooperation with agonists of different TLRs induced significantly better activation of APCs compared to the effect of TLRs alone. Thus, the GA extract expresses better immunostimulatory potential in cooperation with TLR4 agonist in up-regulation of PMs functional characteristics compared to GC extract as well as induction of better allostimulatory and Th1 polarization capacity of MoDCs treated in cooperation with TLR3 and TLR7 antagonists. Therefore, the extract of alternatively obtained basidiocarps could be an effective additional agent during simultaneous engagements of different TLRs on APCs for *in vitro* preparation of APCs as a tool for anti-tumor therapy.

Today, it is well known that fungi possess the potential for production of high diversity low molecular weight products (secondary metabolites) with various biological activities that are mainly important for their interactions with other organisms (Brakhage and Schroeckh 2011). These secondary metabolites are present in a small amount and their composition/combination depends on the type of substrate on which they are cultivated. It may be a reason for different immunomodulatory effects of GC and GA extracts noted in this study. The literature data and data of this study indicate that the study of the *in vitro* cultivation conditions that could possibly optimize their production seems of major importance. Another very important fact in the recent modern period is the need to find a cost-effective and environmentally friendly alternative substrate for *G. lucidum* cultivation which classically was grown on sawdust of sheesham, mango, and poplar.

The genus Ganoderma (especially *G. lucidum),* has been used since ancient times in Eastern traditional medicine. In recent years, the precise effect and their mechanisms were investigated and *G. lucidum* in modern days is used in the treatment and prevention of various pathological conditions. The numerous studies have already demonstrated that various *Ganoderma* spp. extracts and metabolites possess the strong immunostimulatory activity and present effective modifiers of some biological response (Lin, Chen et al. 2006, Boh 2013, Shi, Zhang et al. 2013, Liu, Xing et al. 2016, Liu, Xing et al. 2016). Also, Pi et al. (Pi, Chu et al. 2014) and Lin et al. (Lin, Chen et al. 2006) noted remarkable activation of Th1 and Th2 cells and certain cytokines synthesis in mice treated with *G. tsugae* hot water extract and *G. formosanum* polysaccharide fraction. Similarly, *G. lucidum* polysaccharide, besides effective stimulation of Th1/Th2 immune response, caused stronger proliferation of murine macrophages and significantly higher synthesis of NO as well as IFN-γ, TNF-α, IL-4 and IL-6 in comparison with the control group (Shi, Zhang et al. 2013, Liu, Xing et al. 2016, Liu, Xing et al. 2016).

Regarding the crucial role of macrophages during establishing and maintaining homeostasis and defending against pathogens and transformed cells, these cells are involved in pathogenesis in many diseases (Jung, Kim et al. 2015, Ampem, Azegrouz et al. 2016, Da Silva and Barton 2016). In response to differences in soluble characteristics of a microenvironment and different signals from microorganism-associated molecular patterns, macrophages can polarize into pro-inflammatory, M1, or anti-inflammatory, M2, phenotype (Lawrence and Natoli 2011, Murray and Wynn 2011). *In vitro* cultivation of PMs from mice/rats represents an exceptionally powerful technique to investigate macrophage functions in response to different stimuli, resembling as much as possible the conditions observed in various pathophysiological conditions or as potential therapeutically agents. Thus, glucan isolated from *G. lucidum* spores has also stimulated cytokine production by PMs in culture (Guo, Xie et al. 2009). The considerable immunostimulatory effect was also exhibited by *G. atrum* polysaccharide as well as its acetylated form, which caused increased viability of T lymphocytes and level of IL-2 and TNF-α in the serum of immunosuppressive mice treated with cyclophosphamide overdoses, by regulation of ROS production and NF-κB activity (Chen, Zhang et al. 2014, Yu, Nie et al. 2014, Li, Li et al. 2017). In this study, the immunomodulatory potential of GC and GA extracts was evaluated on PMs with or without TLR4 cooperation (LPS). Macrophages stimulated with LPS are termed as classical activation macrophages (M1 macrophages) and they are involved in the inflammation, pathogen clearance, and anti-tumor immunity (Atri and Guerfali 2018, Shapouri-Moghaddam, Mohammadian et al. 2018). Results of this study show that GC extract with LPS as TLR4 agonist expressed potential for stimulation of metabolic activity/viability of PMs and stimulation of NO production by PMs, while do not express the potential for modulation of adhesive and phagocytic potential and ROS production by PMs. On the other hand, GA extract in combination with TLR4 signals induced by LPS induces intensive stimulation of all investigated functions of PMs including metabolic activity, phagocytic capacity, and production of ROS and NO compared to the effect of GC. Additionally, GA increased the adhesive capacity of PMA-stimulated PMs. These results may indicate that GA extract with TLR4 agonist induces stronger signalling that is responsible for stimulation of very significant characteristics of M1 macrophages such as their metabolic activity, phagocytic activity, ROS and NO production compared to TLR4 signalling alone. Also, adhesive capacity of PMA-stimulated PMs was additionally increased by GA extracts. A few studies demonstrated the mechanism of mushroom polysaccharides action on cytokine production. Namely, Kim et al. (Kim et al. 2012) and Pi et al. (Pi, Chu et al. 2014)showed that augmentation of TNF-α synthesis by macrophages was based on the polysaccharide binding for TLR4 sited on macrophage membrane and it recognition as pathogen-associated molecular patterns. However, detailed analyses demonstrated that induction of mRNA expression in Sarcoma 180-bearing mice is the main mechanism of the polysaccharide action (Zhang, Nie et al. 2014). Since these studies demonstrated that intracellularly generated ROS (as a response to pathological stimuli) affect NF-κB activation and in such way cytokine production by macrophages, it can be concluded that ethanol extract of alternatively cultivated *G. lucidum* basidiocarps which significantly increased ROS production in PMs, could be a potent immunostimulatory and anti-tumor agent.

Induction of effective adaptive immune responses dependents on signals from innate immunity especially from the level of maturation of DCs and their characteristics (Constantino, Gomes et al. 2017). In our study, the combination of poly I:C, loxoribine and GC/GA extract induced phenotypic maturation of MoDCs as determined by u-regulating the surface molecules, including HLA-DR and CD83. CD83 acts as a key DC maturation marker (Prechtel, Turza et al. 2007). However, the combination of poly I:C, loxoribine and GA extract induced up-regulation of CD86, CD40 and CCR7. CD86 is a main co-stimulatory ligand for T cells, providing the second signal for proliferation and clone expansion of antigen-specific T cells (Jeannin, Magistrelli et al. 2000). CD40 is also an indicator of the activation state of MoDCs whose up-regulation acts in favour of enhanced T cell activation. Interaction of this molecule with its ligand (CD40L), expressed by activated T cells is important for up-regulation of co-stimulatory molecules on DCs and enhanced capacity of DCs to trigger proliferative responses and for the regulation of DC functions (Ara, Ahmed et al. 2018). It is in line with our results which showed significantly higher proliferation of CD4 T lymphocytes co-cultivated with MoDC treated with the combination of poly I:C, loxoribine and GA extract. It is important to mention that solely maturation of DCs, expressing high levels of co-stimulatory and maturation markers, is not sufficient for an adequate immune response. Namely, various cytokine production and subsequent CD4^+^ T lymphocytes polarization by DCs is of great significance in the induction of proper immune response as well (Zobywalski, Javorovic et al. 2007). Poly I:C is known as a potent stimulator of bioactive IL-12 production and subsequent activator of the Th1 immune response (Rouas, Lewalle et al. 2004). Our findings confirm these published results because Poly I:C in combination with loxoribine and GC/GA extract induces the up-regulation of IL-12 level. An important finding of this study was the intensive promotion of Th1 and slightly promotion of Th17 polarizing capability of MoDCs by simultaneous engagement of poly I:C, loxoribine and GA compared to the capability of MoDCs by simultaneous engagement of poly I:C, loxoribine and GC. Regarding IL-10 production, obtained results are also interesting. Namely, IL-10 is an immunoregulatory cytokine, responsible for tolerogenic properties of DCs (Smits, Engering et al. 2005), which participates in balancing of the immune response (Saraiva, Christensen et al. 2009). Production of IL-10 could be relevant as a down-regulator of extensive production of immunostimulatory cytokines, knowing that the balance between stimulatory and inhibitory cytokines is important for critical point during the immune response. The decreased level of IL-10, in co-culture of CD4^+^ T lymphocytes and MoDCs, treated simultaneous with poly I:C and loxoribine and GC/GA, could be explained by mutual antagonistic effects of Th1 cells on Th2 cells and Tregs as showed by Glimcher and Murphy (Glimcher and Murphy 2000). Therefore, down-regulation of IL-10 production may serve as an additional mechanism for the promotion of the Th1 immune response.

Bearing in mind the significance of anti-tumor vaccines, this study performed to find the optimal protocol for the development of DCs able to induce an adequate immune response. The results that we obtained in this study suggest that simultaneous TLR3 and TLR7 signaling with signaling induced by components of crude extract of *G. lucidum* cultivated on wheat straw may provide a previously undescribed approach for DC-based vaccine development by using synergistic TLR ligands and *G. lucidum* extract. Overall, these results point to that natural immunomodulators possible mechanism for enhancement of effects of known TLR agonists.

## Conclusions

The significance of our results is highlighted by the following facts: (*i*) immunomodulatory activity is known only for wild and traditionally cultivated *G. lucidum* basidiocarps; (*ii*) traditional cultivation of *G. lucidum* basidiocarps on various hardwood sawdusts is not ecologically and economically friendly; (*iii*) substratecomposition and cultivation conditions significantly affected type and content of bioactive metabolites and their activities (Cilerdzic, Sofrenic et al. 2018); (*iv*) wheat straw induced a synthesis of numerous bioactive molecules, primarily triterpenoids (Cilerdzic, Sofrenic et al. 2018), which could be considered as the one of strong modulators of APCs activity; (*v*) there is increasing need for natural immunomodulators, without any side effect; (*vi*) modern vaccines based on highly purified antigen induce insufficient immune protection and a need for natural vaccine adjuvants is continuously growing.

## Acknowledgments

This work was supported by Ministry of Education, Science and Technological Development of the Republic of Serbia (451-03-68/2020-14/ 200178) and Ministry of Defence of the Republic of Serbia (MFVMA/10/13-15).

